# Temporal profiling of redox-dependent heterogeneity in single cells

**DOI:** 10.1101/198408

**Authors:** Meytal Radzinski, Rosi Fasler, Ohad Yogev, William Breuer, Nadav Shai, Jenia Gutin, Tommer Ravid, Nir Friedman, Maya Schuldiner, Dana Reichmann

## Abstract

Cellular redox status affects diverse cellular functions, including proliferation, protein homeostasis, and aging. Thus, individual differences in redox status can give rise to distinct sub-populations even among cells with identical genetic backgrounds. Here, we have created a novel methodology to track redox status at single cell resolution using the redox-sensitive probe roGFP. Our method allows identification and sorting of sub-populations with different oxidation levels in either the cytosol, mitochondria or peroxisomes. Using this approach we defined redox-dependent heterogeneity of yeast cells, and characterized growth, as well as proteomic and transcriptomic profiles of subpopulations of cells that differ in their redox status, but are similar in age. We report that, starting in late logarithmic growth, cells of the same age have a bi-modal distribution of oxidation status. A comparative proteomic analysis between these populations identified three key proteins, Hsp30, Dhh1, and Pnc1, which affect basal oxidation levels and may serve as first line of defense proteins in redox homeostasis.

## Introduction

Cellular redox status has long been known to play a role in aging and cell functions, with increasing evidence linking aging-related diseases to changes in oxidation levels in a variety of organisms ^1-3^. Oxidative stress has been found to correlate with the generation of reactive oxygen species (ROS), protein aggregation, and apoptosis^4,5^, as well as a number of other physiological changes that ultimately occur in a heterogenic manner.

A series of useful probes developed in recent years have enabled assessment of redox changes within the cell^6-8^. These are applicable in multiple organisms and can be targeted to sub-cellular compartments to report changes within specific organelles. While some of the probes are non-optical or luminescence-based, the majority are fluorescence-based, meant to identify changes in either signaling and regulatory oxidants, or redox homeostasis related redox couples such as glutathione (GSH/GSSG). One prominent redox sensor is the cytoplasmic redox-sensitive GFP variant (roGFP) fused to Glutathione 1 (Grx1) known as Grx1-roGFP2, with additional variants targeting it to organelles such as the mitochondria or peroxisome. Other roGFP-based sensors specifically monitor changes in hydrogen peroxide (H_2_O_2_) levels, through use of thioredoxin peroxidases fused to roGFP2 molecules. These and other similar probes have been used to conduct real-time measurements of redox potentials in bacteria^9^, *C. elegans*^10^, and plant^11^ and mammalian cells^12^, by monitoring differences in oxidative status under a range of diverse conditions.

Detection of roGFP redox-dependent fluorescence has generally been based either on imaging individual cells by microscopy, or by measuring the total fluorescence signals of cells in suspension by using plate readers. However, neither approach enables high spatiotemporal resolution in widescale tracking cell to cell diversity, nor subsequent isolation of cells based on their redox status. Over the last decade, numerous studies have pointed to the fact that populations of genetically identical cells are heterogeneous in their protein and gene expression^13,14^, exhibiting an array of differences in cellular behavior and in varying abilities to respond to changing environments^15-17^. This cell-to-cell variability is considered to be one of the crucial features in the evolution of new survival strategies in fluctuating environments^16^, antibiotic treatment^18^, pathogen progression^19,20^ and other processes. However, the cell-to-cell heterogeneity of redox status within a population of synchronized cells with an identical genetic background has not yet been explored.

Here, we developed a highly sensitive methodology based on the Grx1-fused roGFP2 redox sensor that uses flow cytometry to measure the redox state of individual cells within a heterogeneous *saccharomyces cerevisiae* (henceforth referred to as yeast) population during chronological aging. Sorting of the yeast cells based on their oxidation status allowed us to define the phenotypic, proteomic and transcriptomic profiles associated with the redox state of genetically identical cells of similar age. We show that the proteome and transcriptome profiles of reduced and oxidized cells differ within a yeast population, in addition to corresponding changes in growth and cellular division. Comparative proteomic analysis identified three key proteins: the chaperone Hsp30, the helicase Dhh1, and the nicotinamidase Pnc1, which affect basal oxidation levels and might serve as first line of defense proteins in redox homeostasis.

We also demonstrate that although the ratio between the oxidized and reduced yeast subpopulations changes during chronological aging, the major features, including the transcriptome and proteome, remain relatively unchanged during their first two days. By using cell imaging, we further show that there is a threshold of oxidation, above which the cell cannot maintain redox homeostasis. Finally, microscopic observations of budding cells show that once a mother cell is close to or above this threshold, it passes the oxidized state onto the daughter cell, which starts its life from a high, inherited oxidation level.

## Results

### Flow cytometry based methodology provides a highly accurate way to monitor the subcellular redox status of individual yeast cells

Cellular redox status has been suggested to be correlated with cell function and longevity^21^. Measurements of cellular oxidation levels tend to be based on a global assessment of protein oxidation and the cellular redox status of the entire cellular population^22^. Intrigued by the possible heterogeneity of wild type yeast cells' oxidation levels, we developed a highly sensitive flow cytometry-based methodology to monitor and quantify the cellular redox status and sort out subpopulations of cells based on their oxidation levels.

To do so, we utilized the redox sensitive GFP variant Grx1-roGFP2 (Fig. 1A)^23^, which has a characteristic fluorescence according to a redox-dependent conformational change^24^ that leads to an alternating peak intensity at 405nm and 488nm. In order to verify roGFP expression patterns, we measured the fluorescence spectrum of cells between 360nm and 495nm, following emission at 535nm, under either oxidizing or reducing conditions (Fig. 1B). In agreement with previous studies, we identified the alternating peak pattern at 405nm and 488nm^24^.

**Figure 1.**
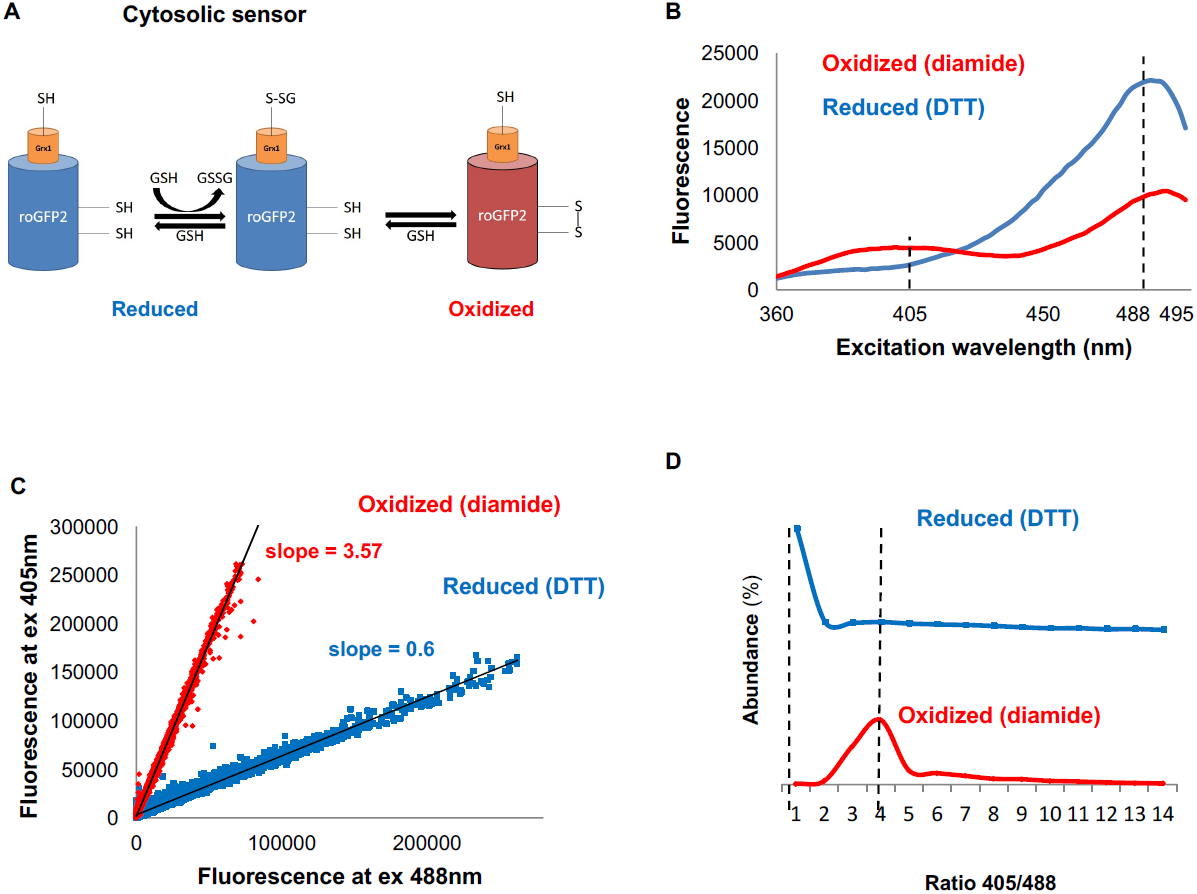
Monitoring subcellular redox levels in yeast cells using the Grx1-roGFP variants and flow cytometry. (A) Scheme of the Grx1-roGFP variant used in this study to monitor oxidation in the cytosol. Cellular GSSG reacts with the catalytic residues of the fused Grx1 which leads to oxidation of the modified GFP protein. (B) Fluorescence excitation spectra of Grx1-roGFP in fully reduced (blue) and fully oxidized (red) yeast cells. Emission followed at 510nm. (C-D) Quantification of redox status of fully reduced (blue) and fully oxidized (red) cells using FACS. (C) Fluorescence of Grx1-roGFPat 510nm obtained using excitation by 405mn and 488nm lasers. (D) Distribution of the 405nm/488nm ratios among fully reduced and fully oxidized cells.

To dissect cell-to-cell variation, we applied flow cytometry to quantify the ratiometric fluorescence intensity of individual cells in a synchronized population at 405nm and 488nm. First, we determined the degree of cellular oxidation by measuring the differential roGFP fluorescence under oxidative and reducing conditions (15 minute treatment with 8mM Diamide and 40mM DTT, respectively). To focus on living cells we gated out the dead and roGFP negative cells, leaving only cells that had a fluorescence intensity at both 405nm and 488nm. The advantage of using the ratiometric approach is that the outcome does not depend on probe expression, which can vary with age and/or treatments. The 405/488nm ratios of reduced and oxidized cells were significantly different and could be characterized using linear fits with a slope of 3.57 for reduced and 0.6 for oxidized cells (the correlation coefficients (R^2^) were 0.99 and 0.97 for oxidized and reduced cells, respectively) (Fig. 1C, D). This approach enabled us to quantitatively determine a normalized value (OxD), as described in the methods section, and to apply it to the quantification of the relative redox status of the cells in yeast cultures.

The two-fold difference between the oxidized and reduced states enabled us to create gates for oxidized and reduced cells (Fig. S1), such that we could sort yeast subpopulations according to their redox status. This approach was also applied to organelle-specific redox sensors, such as the mitochondrial Grx1-roGFP2-Su9^22^ and the peroxisomal Grx1-roGFP2-SKL^25^, revealing a similar redox sensitivity (Fig. S1).

### Utilizing a dual sensing system to define a redox profile in the cytosol and peroxisomes

To examine whether flow cytometry was accurate enough to capture even changes at the subcellular level, we applied our method to examine the redox crosstalk between peroxisomes and the cytosol. To do so, we expressed either the cytoplasmic or peroxisomal (Peroxisomal Targeting Signal type 1 at sensor end: Grx1-roGFP-SKL)^25^ roGFP sensor (S2A) in the background of four peroxisomal gene deletions, each designed to report on a different aspect of the probe: *Δpex3*, *Δcat2*, *Δatg36* and *Δahp1*. Using flow cytometry, we monitored OxD levels in the wild type and deletion strains at the late-log phase (Fig. 2, supplementary table 1). In general, oxidation levels in the peroxisome were slightly increased relative to the cytosol (Fig. 2A, F). As a control we deleted Pex3, an essential peroxisomal protein^26^, leading to a complete loss of peroxisomes and the distribution of the peroxisomal sensor back to the cytosol (Fig. 2B, F). Since the roGFP-SKL sensor becomes cytosolic, we could indeed measure that both probes gave near-identical results, with a small decrease in the OxD values for the peroxisomal roGFP sensor (Fig. 2B). A difference in the OxD level of the peroxisomal roGFP sensor between wild type and Pex3 null cells suggested that, as expected, the peroxisome is slightly more oxidizing under normal conditions. Deletion of the peroxisomal carnitine acetyl-CoA transferase (Cat2) resulted in slightly oxidized peroxisomes with no significant effect on the cytosolic redox status (Fig. 2C, F, supplementary table 1). This is consistent with the finding that deletion of Cat2 leads to sensitivity to oxidative stress^27^. Deletion of Atg36 - a required protein for peroxisomal autophagy - had no effect on the cytoplasmic OxD, however it reduced peroxisomal redox levels two-fold (Fig. 2D, F), suggesting that in the absence of pexophagy, malfunctioning peroxisomes accumulate and hence have a similar redox status as the cytosol. Another deletion that led to slightly increased oxidation was the deletion of a thiol-specific peroxiredoxin, Ahp1, which reduces peroxide levels. Subcellular localization of Ahp1 is not fully defined and it has been detected in the cytosol, however it also has a peroxisomal signal peptide, suggesting a peroxisomal localization^28,29^. Moreover, the *Candida boidinii* orthologous protein is localized to the peroxisome. Based on our results, we speculate that Ahp1 plays an important role in maintaining redox status in the peroxisome in addition to the cytosol.

**Figure 2.**
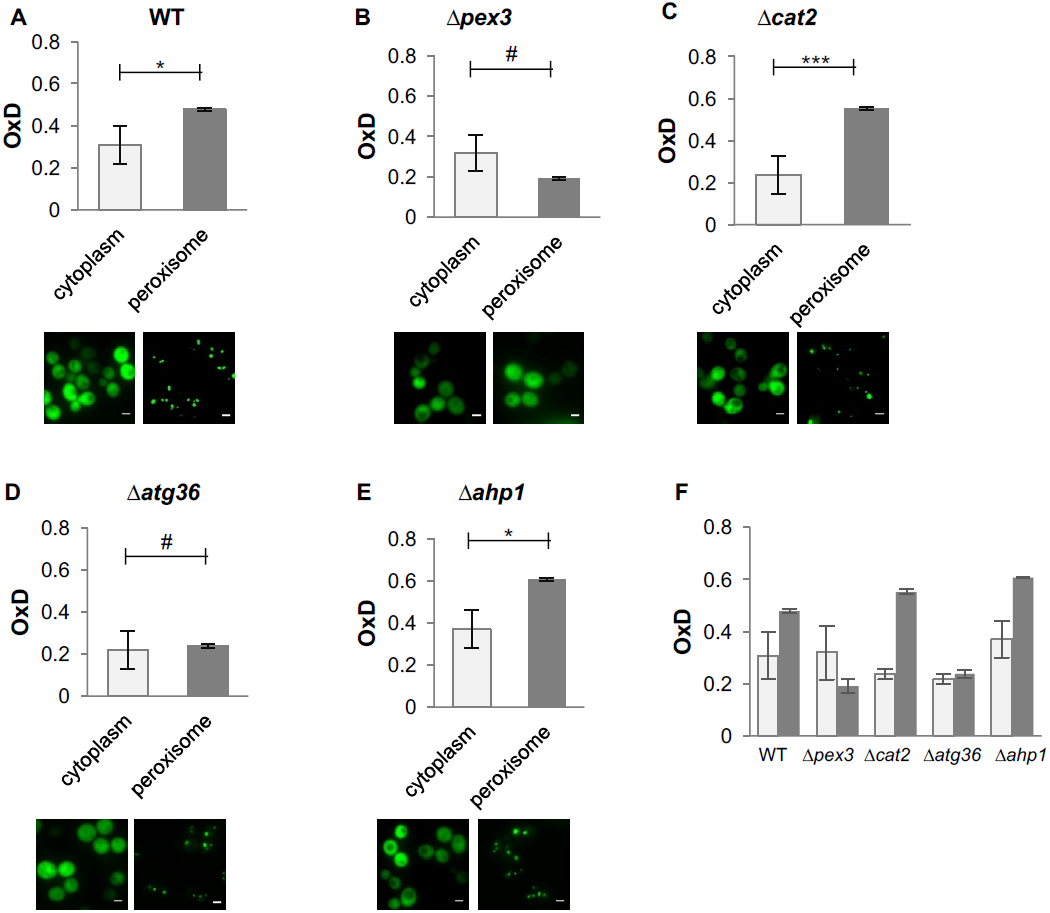
Flow cytometry can detect both cytosolic and peroxisomal redox changes. Cells expressing the cytoplasmic roGFP sensor or peroxisomal roGFP in wild type (A) or in the background of different peroxisomal gene deletions (BE) were grown for 24h and their OxD measured. (F) The same measurements as in A-E, summarized in the same plot. Representative images of the roGFP signal for each organelle are shown in Fig. S2. The flow cytometry method is sensitive such that it can accurately capture the oxidation changes that occur not only in the cytosol but also in the peroxisome itself.

Overall, these findings demonstrate the sensitivity of our approach and its ability to detect even slight changes in anti-oxidant activity within even the smallest of organelles. Moreover, it indicates the robustness of cytosolic redox maintenance mechanisms, despite changes in peroxisomal oxidation (Fig. 2F).

### Defining the redox-dependent heterogeneity in an aging population

The association between chronological aging and cellular oxidation has been explored in previous studies, with research indicating that older cells in a culture undergo changes in their oxidative stress response^30,31^. To examine the ability of our flow-cytometry based method to monitor the age-related oxidation in yeast cells, we first used the cytosolic Grx1-roGFP2 redox sensor to evaluate the average cellular redox changes of the wild type and *Δglr1* strain (which lacks the glutathione reductase gene) over four days of growth in standard medium enriched with casein digest. Since glutathione reductase reduces oxidized glutathione and mainly maintains the glutathione state in the mitochondrial intermembrane space^32^, we predicted that the null mutant would have higher cytosolic and mitochondrial oxidation. As expected, cellular oxidation increased with increasing chronological age in both strains, however the *Δglr1* mutant had relatively higher OxD levels as compared to wild type (Fig. 3A, black circles). Interestingly, when using the mitochondrial sensor (Grx1-roGFP2-Su9) on the same wild type strain, we identified a smaller age-dependent difference in mitochondrial versus cytosolic oxidation, suggesting a robustness of the mitochondrial system's antioxidant behavior during chronological aging (Fig. 3B, white circles). This result is consistent with previously published studies, demonstrating that lower mitochondrial oxidation is maintained throughout chronological aging^22^.

**Figure 3.**
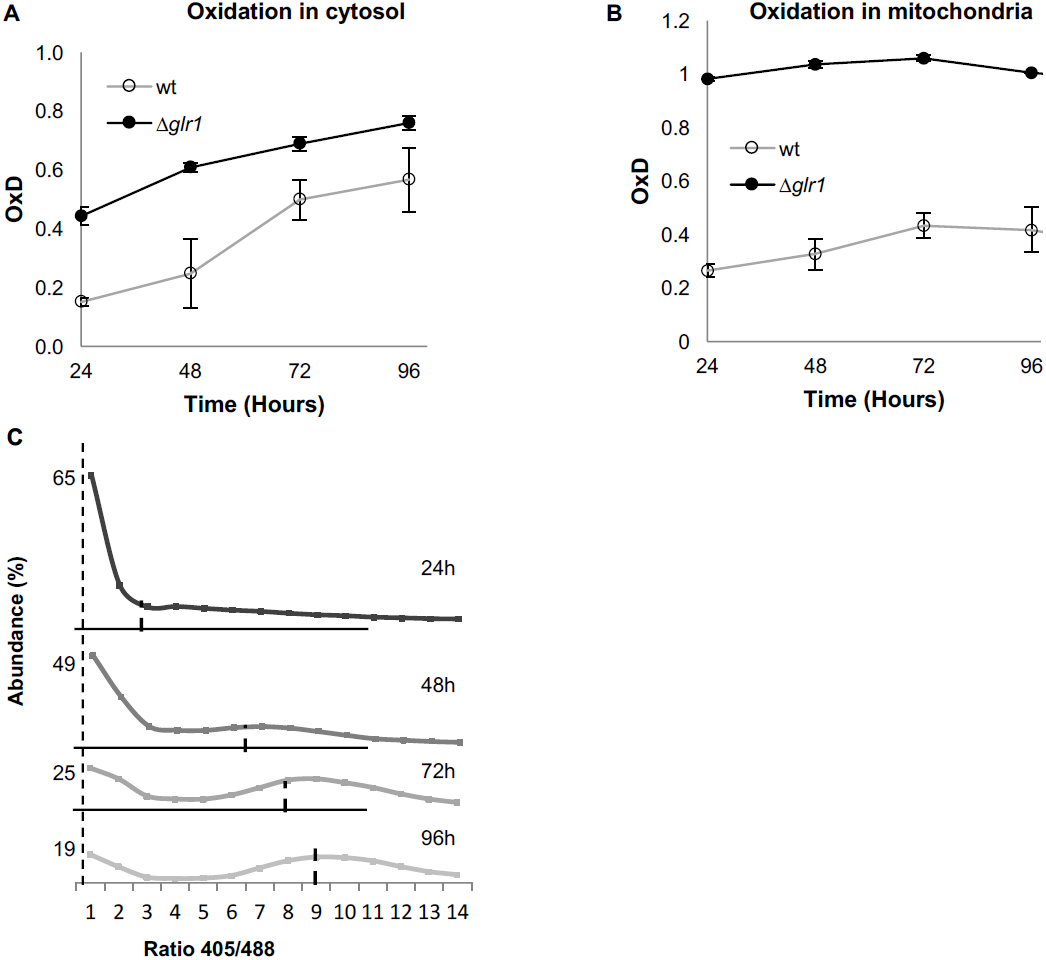
Both cytosolic and mitochondrial oxidation levels increase with age and indicate the emergence of oxidized and reduced cellular subpopulations. Cells expressing cytosolic Grx1-roGFP (A) or mitochondrial Grx1-roGFP-Su9 (B) probes in wild type cells or in the background of a glr1 deletion were grown for four days and their oxidation level monitored. In each experiment 10,000 cells were measured. Shown are oxidation changes over four days in both the cytosol and mitochondria. (C) Bi-modal distribution of ratios of fluorescence intensities obtained at 405nm and 488nm in yeast samples of different ages (24h, 48h, 72h, and 96h). A peak at ratio ≤ 1 represents the reduced subpopulation, while ≥ 6 represents the oxidized subpopulation.

Furthermore, knockout of the *glr1* gene resulted in relatively high and constant mitochondrial oxidation over four days (Fig. 3B, black circles vs white circles) due to failure in reduction of oxidized mitochondrial glutathione. Differences in the redox profiles between cytosolic and mitochondrial oxidation underscore Glr1's role in maintaining redox status in the mitochondria and cytosol during chronological aging.

Although the average cytosolic OxD clearly shows an age-dependent increase in oxidation, it does not give an indication of the potential heterogeneity among cells of the same age. When we examined the distribution of the fluorescence intensity ratios of Grx1-roGFP (at 405nm and 488nm) over four days of growth under chronological aging conditions (24h, 48h, 72h, 96h), we found a clear bi-modal distribution of reduced and oxidized subpopulations (Fig. 3C). As the culture aged, the fraction of reduced cells decreased, while the fraction of oxidized cells increased. Notably, the largest shift to oxidation occurred after 48 hours growth. Thus, we were able to monitor the time-dependent shift from reduced to oxidized subpopulations as the yeast cells aged.

### Age-independent growth differences between the reduced and oxidized subpopulations

As our original expectation, led by the current view of the field, was of a gradual increase in the cellular oxidation status of all cells, we were intrigued by the observed bi-modal distribution. Therefore, we decided to characterize in greater detail the reduced and oxidized subpopulations of aging cells. We utilized the previously described predefined flow cytometry gates to isolate these subpopulations by fluorescence activated cell sorting (FACS). The sorted cell subpopulations (reduced and oxidized) were divided into three aliquots: to monitor growth rate, quantify division events, and conduct proteomic and transcriptomic analysis. This process was repeated at least three times in order to provide biological replicates. Due to the low number of oxidized cells obtained after 24h growth, we focused on cultures aged 48h, 72h and 96h.

The subpopulations of the same age (48h or 72h) displayed characteristic differences in their recovery growth rate, whereas the reduced cells exhibited a significantly faster minimal doubling time (MDT) at 48h and 72h relative to the oxidized population of the same age (Fig. 4A). These differences are indicative of an apparent impairment in the oxidized population's ability to rapidly divide and grow as a culture, rather than any inherent inability to enter the logarithmic growth stage (as evidenced by the similar lag phase lengths, Fig. 4B).

**Figure 4.**
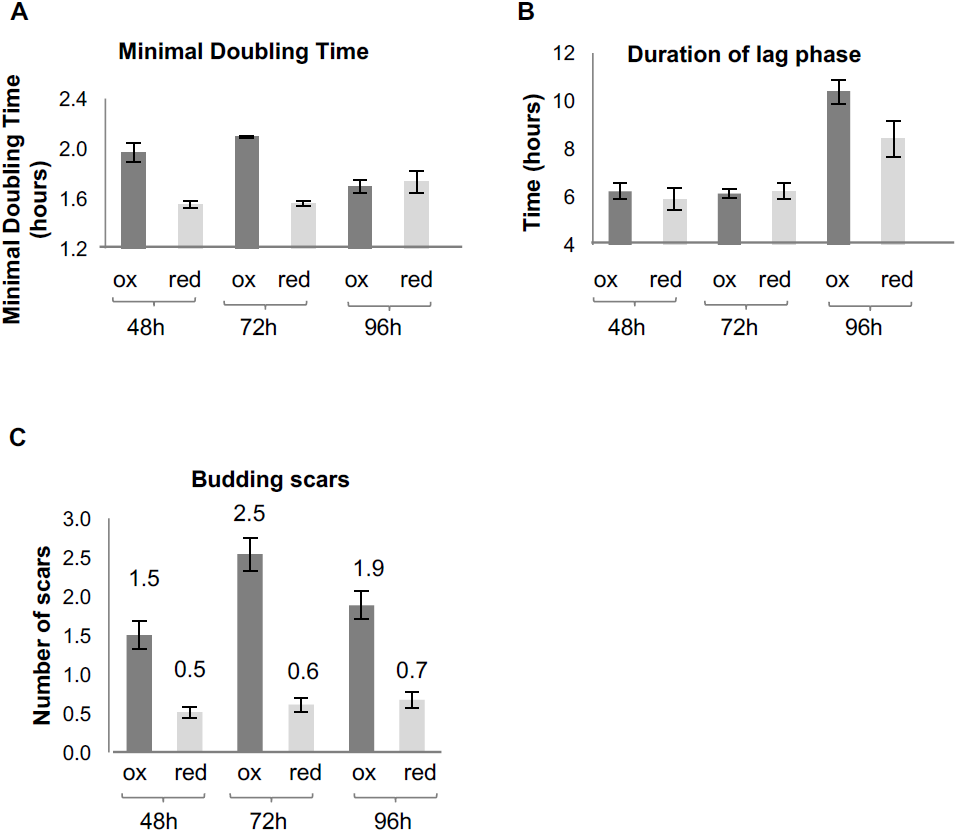
Growth and division of reduced and oxidized subpopulations of yeast cells after sorting by FACS. (A) Differences in minimal doubling time between the sorted oxidized and reduced subpopulations at different time points (48h, 72h, and 96h) measured by OD_600_ in a plate reader. (B) Corresponding duration of the lag phase during recovery growth, measured by plate reader. (C) Corresponding differences in average number of budding scars counted in sorted oxidized and reduced subpopulations assessed by confocal microscopy.

After 96h, both subpopulations had ostensibly similar doubling times following longer lag phases than younger cells (Fig. 4B), consistent with the fact that these were relatively old cells with decreased vitality, particularly among the oxidized cells.

The differences in the growth rates were further supported by a difference in the average number of division or bud scars per cell (Fig. 4C). The oxidized population displayed on average 3-4.5 times more division scars than the reduced population, which typically displayed cells with no more than one scar (Fig. 4C), indicating that the oxidized subpopulation had undergone multiple divisions in contrast to the reduced subpopulation.

Furthermore, the oxidized subpopulation displayed a far lower budding rate relative to the reduced subpopulation, in which approximately 30% of cells at 48h were found to be in the process of budding (Table 1). These results indicate a correlation between the degree of cellular oxidation, growth, and division events. Taken together, the growth and division analysis showed that the differential properties of the subpopulations are redox-, but not age-dependent, at least through 72h growth in standard medium. Rather, the redox state correlates with an intrinsic propensity of cells to divide during the stationary phase.

**Table 1.** Number of budding events

| Growth (h) | Oxidized (%) | Reduced (%) |
| --- | --- | --- |
| 48 | 10 | 30 |
| 72 | 9 | 21 |
| 96 | 6 | 15 |

### Comparison of proteome profiles of the reduced and oxidized subpopulations

In order to further characterize the oxidized and reduced subpopulations of cells, we utilized mass spectrometry to profile the proteome of these subpopulations after 48 and 72 hours of growth. Three biological replicates of each sample group were collected, lysed, trypsin-digested and analyzed by liquid-chromatography-mass spectrometry. Using MaxQuant analysis and a stringent filter, we identified 3389 proteins, of which 1019 were identified in all four subpopulations at least twice (Table S2). According to the Uniprot-based cellular localization annotation, the majority of our identified proteins were cytosolic (51%), where the remaining proteins associated with the mitochondria (16%), nucleus (17%), ER (6%), Golgi vesicles (4%), vacuole (3%), stress granules, P-bodies (0.7%) and peroxisomes (1%) (Table S2, Fig. S3). This is consistent with general distributions of yeast proteins (47% cytosol, 15% mitochondria and 13% ER and secretory vesicles^33^) suggesting that no dramatic expansion or shrinking of an organelle occurred during the experiment.

To obtain further insights into the global changes in the proteome profile of the oxidized and reduced subpopulations, we utilized label-free quantification (LFQ)^34^ to compare expression of the identified proteins between the sample groups. Clustering of average LFQ intensities for all sample groups revealed that the protein expression profiles were similar between groups with a similar redox status, regardless of the sample age (48h or 72h) (Fig. 5A and Fig. S4). The two largest clusters, comprising more than 600 proteins, showed that the differential expression between oxidized and reduced subpopulations remained consistent over two days (Fig. 5A, clusters 3 and 10, Table S2). Annotation enrichment analysis of the proteins comprising these two clusters suggested that the reduced cells (48h and 72h) had an increased presence of proteins involved in energy production, including mitochondrial proteins, protein biogenesis and protein degradation (Fig. 5B). This corresponds with our previous finding that the reduced subpopulation had an elevated growth rate and was in the process of division. The oxidized cells had a distinct subset of proteins regulating protein folding and redox homeostasis, alongside oxidoreductases and NAD binding proteins (Fig. 5B).

**Figure 5.**
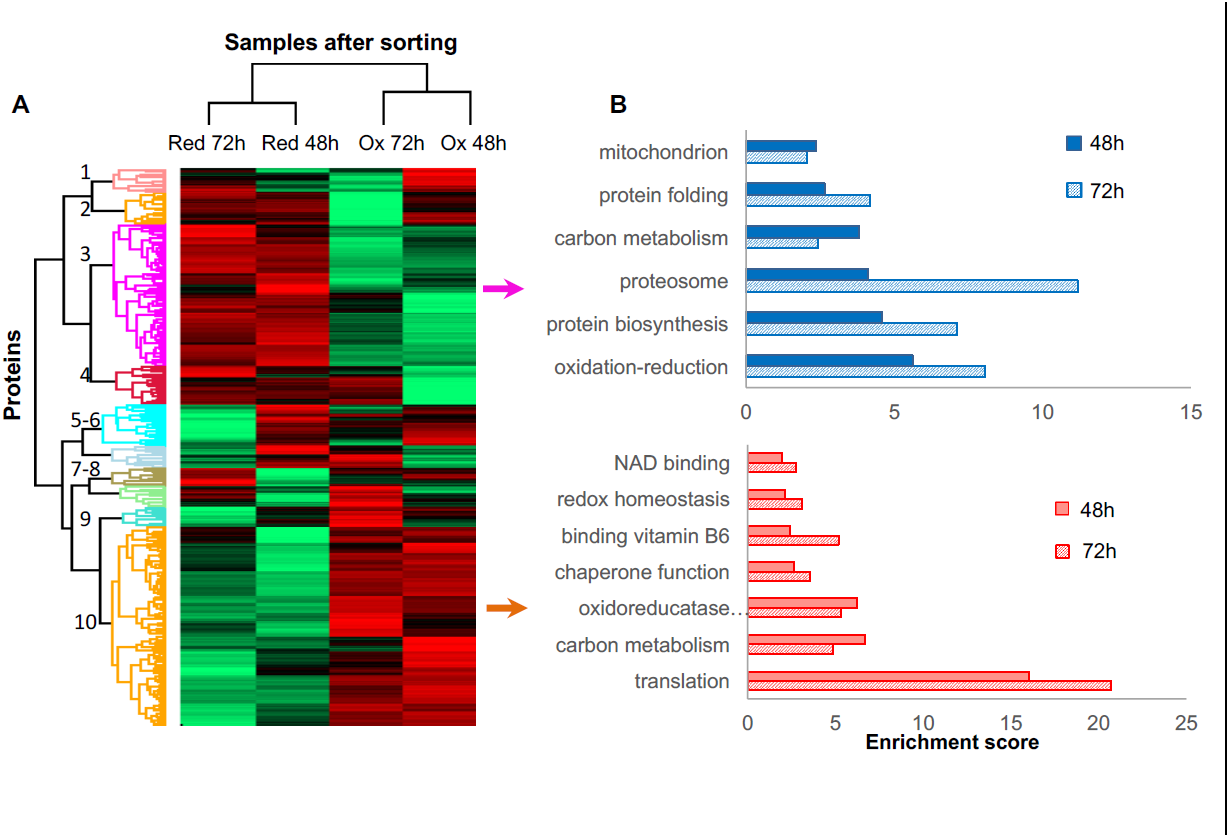
Proteomic analysis and functional enrichment analysis of the reduced and oxidized subpopulations. (A) Hierarchical clustering of proteins identified in cells sorted from cultures at different time points (48h and 72h); downregulated proteins are in green, up-regulated are in red. (B) Functional enrichment analysis of the two largest differentially expressed clusters (3 and 10, representing enriched functions in the reduced and oxidized subpopulations, respectively) at 48h and 72h (solid fill and slashes, respectively).

### Differences in the levels of specific proteins between the reduced and oxidized cell subpopulations

One of the big unanswered questions arising from our data is how do cells of the same age co-exist at different oxidation levels. To identify potential key proteins mediating the redox status of cells, we applied a stringent T-test analysis (false detection rate [FDR] =0.05) to compare protein expression in the oxidized and reduced population of the same age. We identified a subset of 199 proteins showing statistically significant different expression profiles between the reduced and oxidized populations after 48h growth and 149 proteins after 72h growth (Fig. 6A, Tables S3 and S4). Annotation enrichment analysis showed a similar functional classification for clusters 3 and 10 in Fig. 5 (Fig. S5), suggesting that the reduced cells were more metabolically active, expressing higher levels of mitochondrial and carbon metabolism proteins than the oxidized subpopulation. We hypothesized that clustered proteins might belong to a common pathway which would be significantly up- or down-regulated in a redox-dependent manner. Hence, to identify potential protein-protein interactions within the subset of differentially expressed proteins, we used the STRING protein interaction database^35^ to characterize the interaction networks between these proteins. We obtained numerous clusters of interactions between our identified, differentially expressed proteins, associated with protein translation, mitochondrial activity, and stress response (Fig. S6). Each interaction cluster contained proteins upregulated in either oxidized (red circles) or reduced (blue circles) cells. This suggests a complex anti-oxidation mechanism that may differentiate preventative from responsive activity.

**Figure 6.**
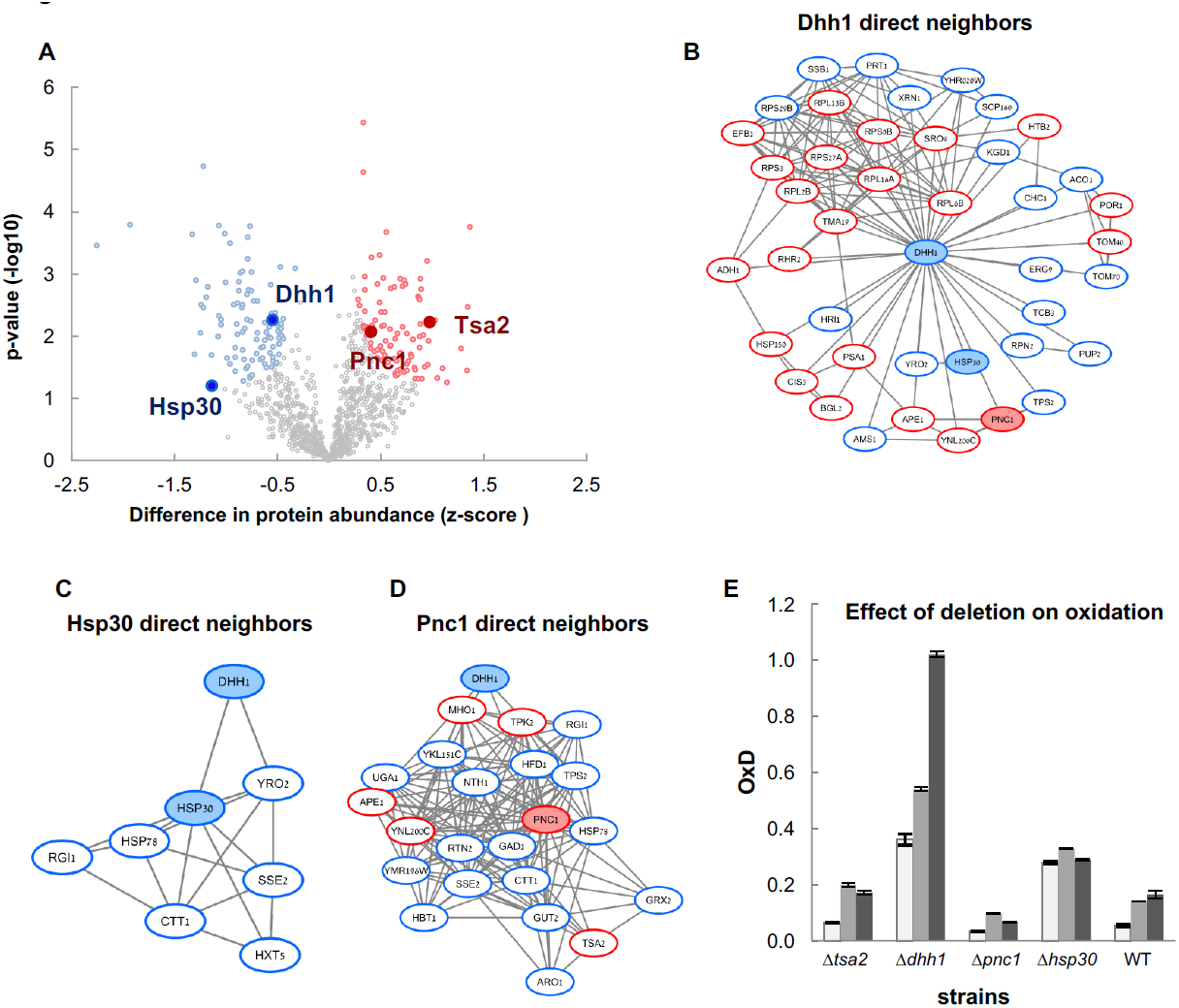
Differentially expressed proteins between the reduced and oxidized subpopulations. (A) Volcano plot of differentially expressed proteins between the reduced and oxidized subpopulations. Significantly expressed proteins are labeled in blue (increased expression in reduced) and red (increased expression in oxidized), according to an FDR of 0.05 and a fold change greater than 2. (B-D) Proteins interaction networks of Dhh11, Hsp30, and Pnc1's direct neighbors based on STRING database interactions of significantly changed proteins (with corresponding colors). (E) Oxidation levels in deletion strains of significantly changed proteins (***Δ****tsa2,* ***Δ****dhh1,* ***Δ****pnc1,* ***Δ****hsp30*, and wild type control) at different ages (24h, 48h, and 72h).

Specifically, the proteome of the oxidized cells contained significant increases in proteins induced by a variety of stresses (Gre1, Sro9, Hsp150), as well as proteins directly involved in redox homeostasis (thioredoxin peroxidase Tsa2, Tma19). Interestingly, the oxidized cells displayed an upregulated expression of ribosomal proteins, in which we identified 21 proteins belonging to 40S and 60S ribosomal subunits, suggesting relatively elevated levels of ribosomes under oxidizing conditions. Conversely, the reduced cells had high levels of proteins related to protein biosynthesis, tRNA-associated proteins, folding factors and chaperones. Hence we speculate that the oxidized cells may accumulate “non-functional” ribosomes whereas the reduced cells were in the process of protein biosynthesis and energy production.

Moreover, elevated levels of cytosolic catalase T (Ctt1) and mitochondrial glutaredoxin (Grx2) were detected in the reduced subpopulation, likely preserving the reduced state of these cells. We also identified the upregulation of several proteins that have been linked to increased longevity, such as Dld1 and Gut2^36^, implying that the redox-dependent expression of these proteins may play an active role in cellular lifespans. Intriguingly, the reduced cells showed increased expression levels of proteins associated with stress granules and stress response, such as stress-induced heat shock proteins: Hsp30, Hsp70-type chaperone Ssb1, its ATP exchange factor, Sse2, disagregase Hsp78, as well as the RNA metabolism proteins: DEAD-box helicase Dhh1, pre-mRNA cleavage protein Hrp1, translational factor Prt1, exonuclease Xrn1 and others (Fig. 6A, Tables S3 and S4). One intriguing possibility is that these proteins serve as a first line of defense against oxidation, such that their expression prevents further cellular oxidation.

### The importance of Hsp30, Dhh1 and Pnc1 in maintaining reduced cellular status

To verify our proteomic analysis and identify potential first line of defense redox proteins, we focused on four selected candidates whose expression significantly increased either in the reduced cells – heat shock protein 30 (Hsp30) and helicase Dhh1 - or in the oxidized cells - thioredoxin peroxidase Tsa2 and nicotinamidase Pnc1 (Fig. 6A). All four proteins have multiple interactions with other significantly differentially expressed proteins identified in our proteomic analysis; Hsp30 is connected to Dhh1 and has few interactions with proteins which are enriched in reduced cells (Fig. 6B, C). Furthermore, Pnc1 and Dhh1 have many known interactions with proteins identified in our analysis and were found to be differentially expressed in reduced and oxidized cells (Fig. 6D).

To examine the contribution of these selected proteins to the cellular redox status over 72h, we measured redox levels using the cytosolic roGFP sensor in the wild type strain, as well as in four knockout strains: *Δtsa2*, *Δdhh1*, *Δpnc1* and *Δhsp30*. Notably, we found that *Δdhh1* and *Δhsp30* were significantly more oxidized than the wild type over 72 hours (Fig. 6E, p-values of the t-test are in Table S6). Both proteins were found to be upregulated in the reduced subpopulation, suggesting that they may play a role in redox regulation pathway mediation. Consistent with previously published studies, the Tsa2 knockout had no significant effect on cellular oxidation^37^ (Fig. 6E Table S6). However, deletion of the nicotinamidase Pnc1, which was found to be upregulated in the oxidized cells, led to a more reduced environment than in wild type cells, specifically after 48 hours of growth (Fig. 6E, Table S6). This corresponds with our proteomic analysis and also suggests that Pnc1 is a potential effector of redox homeostasis.

In general, the comparative analysis of the proteomes for the reduced and oxidized cells at two different ages confirmed the expected redox-dependent divergence with a weak correlation, and identified three potential redox-effectors: Hsp30, Dhh1 and Pnc1.

### A differential transcriptome profile of the reduced and oxidized subpopulations

To examine coupling between protein and transcriptome abundance, we conducted a transcriptomic analysis of three biological replicates of the isolated reduced and oxidized subpopulations at 48h and 72h. We identified 4949 genes in the reduced sample and 5027 genes in the oxidized samples (Table S5).

As expected from the proteomic analysis, global changes in the transcriptome were redox- rather than age- dependent (Fig. 7, and S7). Annotation analysis of the differentially expressed transcripts (defined by an at least two-fold change with FDR<0.05) (Tables S5 and S6) showed relatively similar functional distributions across transcripts from samples of the same age, which correlates well with the annotation of the abundant proteins in these samples. Specifically, the reduced cells had upregulated carbon metabolism and TCA cycle activity. Interestingly, the transcriptomic analysis showed that expression of genes involved in lipid biosynthesis and peroxisomal function were upregulated. This function was missing from the proteomic analysis, most probably due to low total abundance of peroxisomal proteins (˜1%).

**Figure 7.**
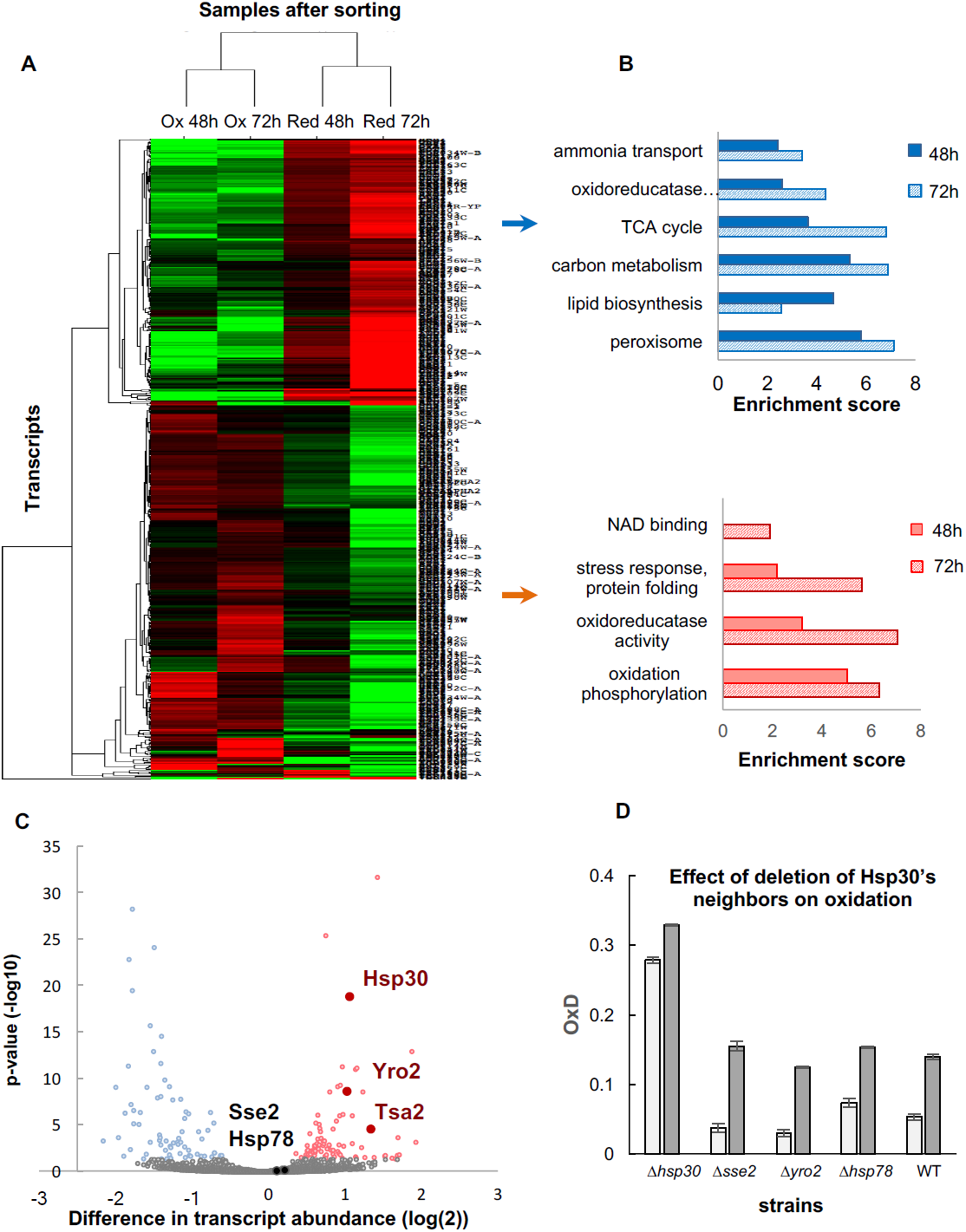
Differentially expressed transcripts between the reduced and oxidized subpopulations. (A) Hierarchical clustering of the median expression values of all differentially expressed genes (FDR<0.05) identified in the post-sorting cells harvested at different time points (48h and 72h); downregulated transcripts are in green, up-regulated are in red. Each row was normalized by its median and the log value was taken for visualization purposes. The data was clustered using a centered correlation similarity metric. (B) Functional enrichment analysis of the differentially expressed transcripts in the reduced (blue) and oxidized (red) subpopulations harvested after 48h (solid bars) and 72h (solid fill and slashes, respectively). (C) The mean value of normalized counts (log2) is plotted for each gene. Each point is colored according to the adjusted value of the differential expression analysis (Methods). Hsp30, Yro2 and Tsa2 were significantly upregulated in the oxidized cells. (D) Oxidation levels in deletion strains of significantly changed orthologues ***Δ****hsp30* and ***Δ****yro2* genes that were not up- or down-regulated; chaperones ***Δ****sse2* and wild type control at different ages (24h and 48h).

Alongside the general agrement between the transcriptomic and proteomic analysis, we found several examples of uncoupling in protein and mRNA expression. One of the intriguing examples is Hsp30 and its homologue, Yro2. Although proteins of these genes were upregulated in the reduced cells, their transcripts were shown to be upregulated in the oxidized cells. As shown previously, Hsp30 is a “redox keeper”; its deletion led to extensive oxidation in cells, correlating with high protein levels in the reduced state. To further investigate the impact of Hsp30 and its associated proteins, Yro2, Hsp78 and Sse2 (Fig. 6) we examined oxidation levels of these protein knockout strains (Fig. 7, Table S11). While Hsp30 had a significant impact on the redox status of cells, as we have already shown, its orthologue Yro2 did not. In addition, chaperones Hsp78 and Sse2, which showed no significant redox-dependent change in their transcript levels, had no influence on the cellular oxidation.

### The transition from reduced to oxidized cellular state is a threshold-based event

To better understand the switch from reduced to oxidized, we tracked redox changes within individual cells in an attempt to identify how cells transition between these two distinct states.

Using confocal microscopy, we monitored OxD changes of specific cells for 12 hours (Fig. 8A). To define the OxD values we measured the fluorescence of fully reduced and fully oxidized cells at 405nm and 488nm (Fig. 8B), as previously described. These measurements were conducted using diluted, unsorted populations of young cells during the early log phase, during which some cells had either lost their plasmid expression or were otherwise roGFP-negative throughout the 12 hours of growth. Nonetheless, we successfully monitored specific OxD changes in over 30 unique cells across 12 hours (Fig 8C-F), and clustered their oxidation trajectories during this time. Interestingly, a subset of these cells maintained their newly reduced state for over 12 hours (the length of the measurement), indicating a strong mechanism for maintaining a reduced environment within the cell (Fig. 8C). An additional portion of these cells lost their GFP signal without an increase in the oxidation-related fluorescence profile. The majority of the reduced cells were able to “self-correct” their oxidative status for relatively prolonged periods (Fig. 8D) or for a shorter time (Fig. 8E) before undergoing oxidation. Surprisingly, this self-correction process existed only below a clear oxidation threshold, at an OxD of approximately 0.7 (Fig. 8C-E). Cells were able to reach OxDs as high as 0.65 before returning to more reduced values as low as 0.4, at which state they could remain for several hours before crossing the threshold and undergoing rapid oxidation. However, we observed that once cells crossed the oxidation threshold, they were unable to correct their oxidative status and maintained high OxD values until their eventual cell death (Fig 8A).

**Figure 8.**
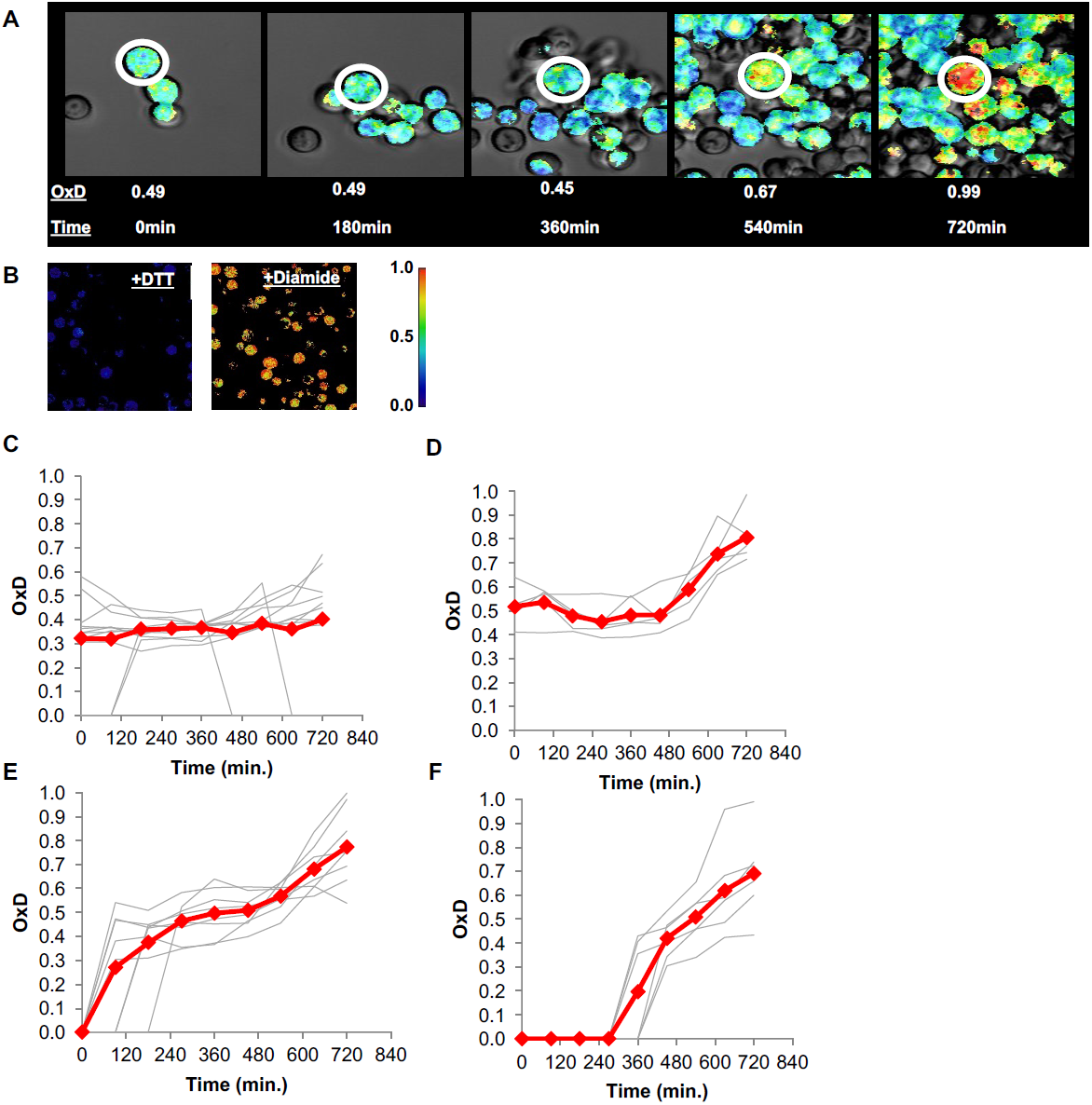
Changes in oxidative status of individual cells over time under confocal microscopy. (A) Oxidation over time of a single cell, from reduced to highly oxidized, imaged using confocal microscopy. (B) Samples treated with 40mM DTT and 8mM Diamide for 15 minutes and imaged using confocal microscopy. (C-F) Oxidation of thirty unique single cells over 12 hours, clustered by trajectory similarity (labeled in red).

Previous research has indicated that in cases of old mother cells with upwards of 10 divisions, the mother cell's age has an impact on its daughters^38^, leading to reduced lifespans in the new daughter cells. To test the suggestion that some division-dependent aging factor was somehow capable of being transferred to the daughter cells, we looked at cases of OxD changes in budding mother cells. We measured the mother cells OxD during and immediately after division, as well as an hour after division. Remarkably, we observed that mother and daughter cells had identical OxD values throughout division and immediately following separation, regardless of the length of division or the mother's initial OxD (Fig. 9), thus demonstrating that daughter cells inherited even a relatively highly oxidized environment during budding. Moreover, some daughter cells were able to self-correct their inherited OxD levels once the budding process was completed and were able to reach a more reduced state within less than an hour, even as the mothers crossed the oxidation threshold or maintained their original OxD (Fig. 9C). These findings suggest that the influence a mother cell has on her daughters may play a more central role in the oxidative stress of the daughter cell in early life.

**Figure 9.**
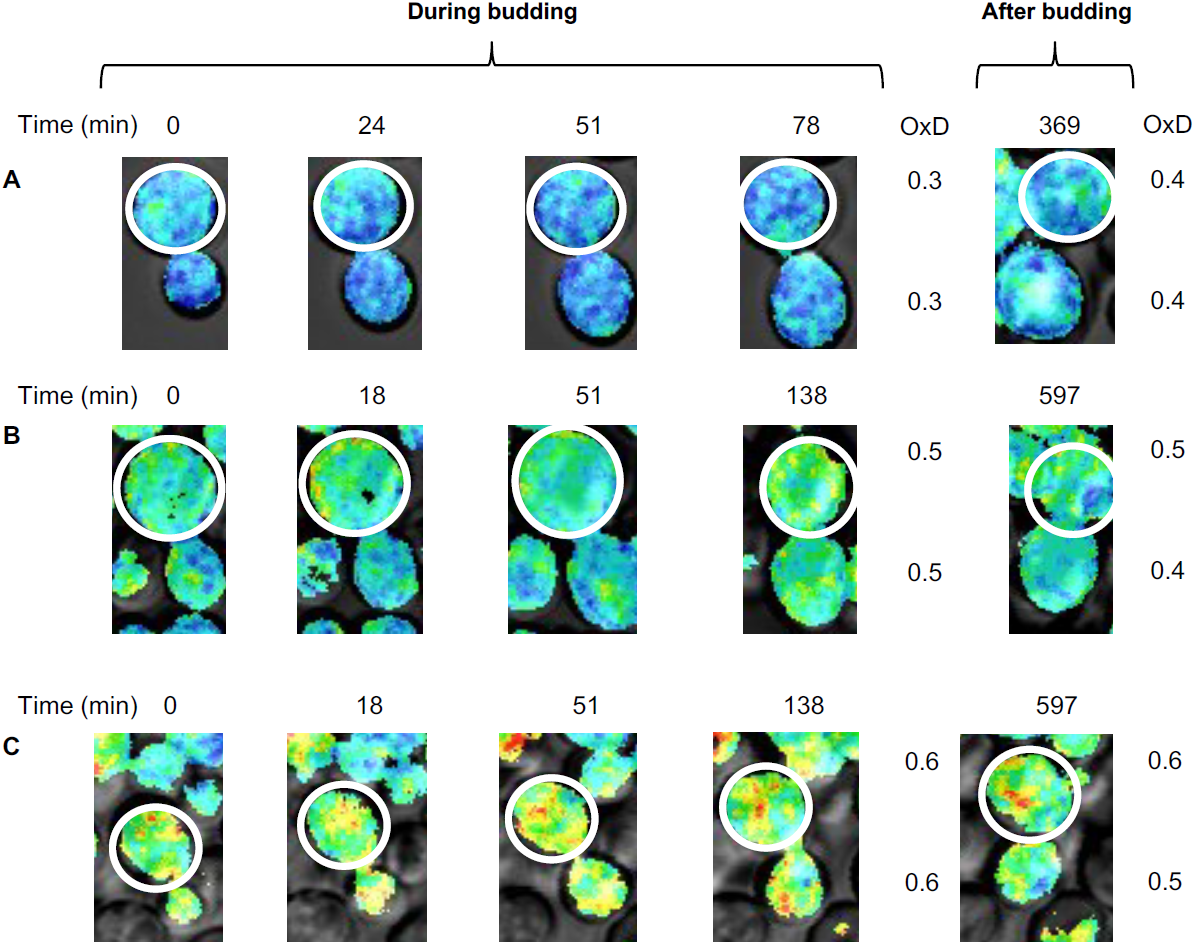
Changes in oxidative status between mother and daughter cells over time under confocal microscopy. (A-C) Oxidation levels over time in mother and respective daughter cells during and after budding, showing the shared oxidative status until separation.

## Discussion

### Dissection of redox-dependent heterogeneity

Phenotypic heterogeneity within genetically homogenous cell populations is considered one of the strategies for dealing with fluctuating environments and stress conditions^13,14,17,39^. Studies in bacteria and yeast have revealed that cell-to-cell variation is encoded in the stochasticity of gene and protein expression in a normal or changing environment. Diverse mechanisms have been suggested, ranging from promoter activation and deactivation, the rate of mRNA and protein production, protein and gene modifications, and even metabolic exchange between cells^40^. The typical readout of these processes is diversity in protein expression leading to differential growth under standard or stress conditions.

Here, we identified a new type of phenotypic heterogeneity, originating in the redox status of cells, by focusing on cytoplasmic oxidation alongside chronological aging. Using a self-established, non-laborious methodology to quantify and isolate cells with different oxidation levels, we suggest that the redox status, rather than cell age, is a critical determinant of cellular growth and division properties. Clearly, other parameters, such as energy and glucose levels, will affect age-dependent oxidation. Nevertheless, on the basis of our monitoring the time-line of cellular oxidation in aging cells in culture, we propose that the transition from reduced to oxidized status is a threshold-based phenomenon that leads to the emergence of at least two distinct cell subpopulations with different growth potential and biochemical profiles.

Our detection of the redox status of aging yeast cells was based on the cytosolic redox-dependent fluorescent probe, Grx1-roGFP2, which depends on GSH/GSSG levels. The roGFP probes are well-established indicators of in-vivo redox potentials in in bacteria^9^, *C. elegans*^10^, yeast^23^, plants^11^, and mammals^12^, monitoring differences in oxidative stress under a range of diverse conditions. Measurements of roGFP fluorescence are generally based on microscopy imaging, quantifying the OxD values for each independently or, conversely, deriving an average OxD value for the entire population of suspended cells. While the first approach is able to evaluate cell-to-cell variability, it is time-consuming and arduous. In the latter approach, direct fluorescence measurement of a population is fast but not sensitive enough to detect natural variations within a defined population. Our utilization of FACS combines the advantages of fluorescence measurement at the level of the individual cell with a high throughput ability to isolate living cells according to their redox-related properties. Due to the robustness of the roGFP sensor, the parameters used to define the oxidation gates to quantify the redox levels in the cytosol, mitochondria and peroxisome were very similar. The discrimination between reduced and oxidized cells was clear-cut, enabling rapid and convenient measurements of biologically different samples collected at different days and conditions. These oxidation gates can be applied to any population of yeast cells expressing the roGFP variants to define the contribution of a specific gene or condition on cytosolic redox status or in organelles. Measurements of this sort may thus also be conducted on a larger scale using knockout-library arrays, ultimately scanning a wide range of mutations and their redox impact across several days. This may contribute to identifying key proteins serving as regulators of redox-dependent heterogeneity or as “redox switches”, defining the ability to respond to various forms of oxidative and environmental stress as well. More generally, this approach is not restricted to yeast and can be used it any cell type - from bacteria to mammals.

### Identification of positive and negative redox modulators: Hsp30, Dhh1, and Pnc1

In this study, we identified three proteins that had a crucial impact on the redox status of yeast at early and late stages of chronological aging: two negative regulators, Hsp30 and Dhh1 that maintain a reducing environment, and one negative regulator, Pnc1, deletion of which decreases cellular oxidation. Hsp30 is a plasma membrane heat shock protein which is induced by different stress conditions, including heat shock, exposure to ethanol, glucose limitation, oxidation and entry into the stationary phase^41^-^44^ During heat shock, deletion of Hsp30 leads to inhibition of the Pma1H+ -ATPase, preserving cellular ATP levels^45^. Moreover, deletion of Hsp30 is correlated with a decrease in superoxide dismutase, *sod1* expression^43^, which might correspond with elevated cellular oxidation. Interestingly, our transcriptomic and proteomic analysis revealed an opposite regulation profile of Hsp30, pointing to an upregulation of gene expression under an oxidizing environment after 48 and 72 hours of growth, alongside downregulation at the protein level at the same time. Moreover, deletion of Hsp30 led to a significant increase in cellular oxidation, suggesting that indeed high Hsp30 protein levels keep the cells reduced. The proteomics analysis showed that upregulation of Hsp30 in the reduced cells was coupled with its native partners, all of which were upregulated in the reduced cells (Fig. 6C: Dhh1, catalase Ctt1, chaperones Hsp78 and Sse2, Hxt5, and Yro2). Interestingly, an orthologue of Hsp30, Yro2, had a similar protein-transcript anti-correlation; however, deletion of Yro2 had a lesser impact on the redox status than Hsp30.

An antagonistic abundance in the RNA and protein levels suggests a potential impairment of Hsp30 protein turnover in oxidized cells. Most probably, gene expression is induced upon oxidation but protein translation is not efficient enough to sustain levels of Hsp30 that would maintain the reducing environment. This correlates with the downregulation of protein biogenesis and energy production, along with increased proteasome activity in the oxidized cells. Thus, enhanced protein turnover (either due to low translation rates or high proteolysis rates) may push the transcriptional machinery to produce high levels of Hsp30 mRNA without reaching critical protein concentration that serves as a “redox keeper” of yeast cells.

In addition to Hsp30, we identified another “redox keeper”, a highly conserved RNA DEAD-box helicase Dhh1, whose protein levels are upregulated in the reduced cells and whose deletion leads to elevated oxidation from early stages of cell growth. Dhh1 is a stress granule-associated helicase, involved in mRNA metabolism^46,47^. Interestingly, other stress granule-associated proteins were upregulated in the reduced cells, including mRNA cleavage factor Hrp1, the translation initiation factor eIF-4B, Tif3, 5'-3' exonuclease, Xrn1, and the eIF3b subunit of the eIF3 translation initiation factor, Prt. This might suggest that downregulation of these proteins leads to a collapse in redox status and migration of these proteins to p-bodies for mRNA decay and transcription inhibition. Interestingly, transcripts of all these proteins, except Hrp1, did not show significant change, suggesting that protein turnover or translation rate might play a significant role in the redox homeostasis, rather than gene expression.

In contrast to the positive effect of Dhh1 and Hsp30, the nicotinamidase Pnc1 was found to be upregulated in the oxidized cells. The null mutant had lower OxD levels than the wild type, suggesting that Pnc1 deficiency in cells with normal redox status might lead to reducing stress, while in oxidized cells its upregulation might have a different effect. Previous studies have shown that deletion of Pnc1 increases nicotinamide levels, which inhibit the histone deacetylase Sir2 and lead to apoptosis or a decrease in life span^48-50^. Thus, we propose that Pnc1 serves as a link between increased reduced stress, nicotinase accumulation, and a decrease in cell viability.

### Threshold-based mechanism and mother-daughter inheritance underline cellular oxidation

Until now it was belived that most oxidative stress comes from age-dependent damage or genetic alterations. However, an intriguing observation of our study is that there were cells with increased OxD values but no division scars in a genetically identical population. To understand the source of oxidation of these “newly born” oxidized cells, we decided to monitor changes in the cellular oxidation utilizing real-time imaging over 12 hours. When examining the distribution of oxidation levels across 12 hours, as expected, we found a clear decrease in the number of reduced cells over time, with the peak shifting towards oxidized, roGFP-negative and damaged cells. Notably, the redox status of cells fluctuated with time, suggesting the activity of a self-repair mechanism until a specific threshold is reached, equivalent to 70% of maximal oxidation; above this threshold, cells remain permanently oxidized.

Analysis of the daughter cells revealed that their oxidation level was identical to that of their mother cells, lasting for several hours “after birth”. Since the oxidized state received by the daughter cells from the mother is maintained for several hours, and thus exceeds their duplication time, we suspect it is a regulated mechanism and not simple diffusion of the roGFP sensor with predefined oxidation. Nevertheless, further research should be done to characterize such redox-related inheritance.

## Methods

### Yeast strains, growth conditions and roGFP probes

The *S. cerevisiae* strains used in this study were the haploid wild-type (BY4741; *MATα, leu2Δ3 his3Δ1 met15Δ0 ura3-0*) and a GLR1 knockout strain (BY4741; *MATα, leu2Δ3 his3Δ1 met15Δ0 ura3-0, GLR1Δ/kanMX*), as well as BY4741 deletion strains made using the pFA6 KO plasmid^51^ provided by the Schuldiner laboratory. The strains were transformed with Grx1-roGFP2, Grx1-Su9-roGFP2, and Grx1-roGFP2-SKL probes^25^ (kindly provided by Bruce Morgan and Maya Schuldiner, respectively) and grown overnight from plate on 2-4ml minimal modified casein amino acid, supplemented with amino acids corresponding to plasmid selection. Cultures were then diluted to an OD_600_ of approximately 0.25 on 4-10ml to ensure fresh growth and brought to their logarithmic phase of OD_600_ of approximately 0.75, which marked “time zero”. Samples were grown at 30°C under constant agitation. Transformations were refreshed every 2-3 weeks in an adapted version of the protocol from^23^.

### Calculation of OxD

OxD (the degree of oxidation) was calculated directly from six measured intensities:

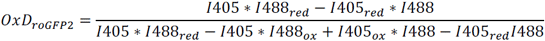

This same equation can be rewritten using direct wavelength ratios:

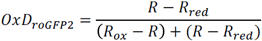

### Monitoring yeast growth

Growth curves were conducted on sorted yeast samples using a TECAN Infinite 200 PRO series, using an adapted version of the protocol from^23^. Measurements were initiated using 4.5 million cells sorted according to their redox status. Yeast incubation and OD_600_ measurements, as well as minimal doubling times, were analyzed using MDTCalc. Minimum doubling time was assessed by first calculating the log10 value of each point during logarithmic growth, and then measuring the slope between each set of consecutive points. The value of this slope was then inverted (representing the minimal time it took the sample point to “double”), providing the minimal doubling time value.

### Flow cytometry analysis of redox ratios and sorting

Yeast cells transformed with roGFP probes and treated with 1:250 (1µl) propidium iodide Sigma p4170 for 15 minutes (adapted from the protocol described by Ocampo at al^52^) were measured during their extended stationary phase every ˜24 hours using no more than 1.0 OD_600_ (10µl), suspended in 240µl phosphate buffered saline (PBS x 1). Total oxidation and reduction of the cultures were determined through addition of 10µl 1M dithiothreitol (DTT) (40mM) and 10µl 0.2M diamide (8mM) to 230µl PBS x 1. Flow cytometry analysis was performed using a sorting-equipped FACS Aria III flow cytometer, with 405nm, 488nm and 531nm lasers (BD Biosciences, San Jose, CA) and the flow cytometry data was analyzed using FACSDiva software (BD Biosciences). Voltage settings for the SSC, FSC, 405nm and GFP channels were kept constant for all experiments. For maximal discrimination between reduced and oxidized roGFP, we used 2D dot plots with linear rather than log scales. Measurements were taken at 405nm and 488nm for OxD calculations and redox ratio, while dead cells were gated out using propidium iodide labeling (excitation at 531nm and emission detected by a 660nm filter). Each analyzed population had a sample size of 10,000 standard cells.

### Sample preparation for mass spectrometry

Sorting was based on the redox ratio, using 9 million cells suspended in PBS. Cells were then lysed using 300µl 0.2M NaOH and resuspended using a lysis buffer (100mM DTT [15.425mg DTT], 100µl 1M Tris HCl pH 7.5, 100µl SDS 20%,complete with DDW to 1ml). Samples were then diluted using 400µl of a urea buffer (8M urea in 0.1M Tris HCl pH 8.5), loaded onto a filter, and centrifuged for 10 minutes at 12,000g following the standard FASP protocol ^53^. This process was repeated three times, with flow-through discarded, after which samples were incubated in the dark for 20-60 minutes with 0.5M iodoacetamide and urea buffer (final iodoacetamide concentration of 0.05M), under constant agitation (350rpm, 25°C). Samples were then again washed three times with urea buffer. The proteins were then digested by use of a digestion buffer (10% ACN, 25mM Tris HCl pH 8.5) and centrifugation at 12,000g for 8 minutes. This step was repeated twice more, after which the filters were transferred to a new collection tube, suspended in 300µl of digestion buffer and 1µl trypsin promega, mixed for 1 minute at 600rpm, and left overnight at 350rmp, 37°. Following digestion, samples were centrifuged for 10 minutes at 12,000g and resuspended in the digestion buffer.

The peptide concentration was determined, after which the peptides were loaded onto stage tips in equal amounts. Stage tips were activated by use of 100µl MS-grade methanol (100% MeOH) and centrifuged for 2 minutes at 2,000g, after which they were cleaned with 100µl of elution buffer (80% ACN, 0.1% formic acid) and centrifuged again for 2 minutes at 2,000g. The stage tips were returned to their hydrophilic state by suspension in 100µl of buffer A (0.1% HPLC-grade TFA) and centrifugation at 2,000g for 2 minutes, repeated once. 10-30µg of protein were then loaded per stage tip (as per protein preparation above), and centrifuged at 1,000g for 2 minutes. Proteins were then washed twice with 100µl buffer A at 1,000g for 2 minutes and transferred to a new collection tube. Peptides were eluted using 60µl buffer B (80% ACN, 0.1% HPLC-grade TFA) centrifuged at 250g for 2 minutes, and 30µl buffer B centrifuged at 250g for 2 minutes. Samples were then dried using a SpeedVac for 18 minutes at 1,300rpm at 35°, after which they were dissolved in 6-12µl of buffer A and prepared for tandem mass spectrometry analysis.

### Nano-LC-MS/MS analysis

The peptides were injected into a Nano Trap Column, 100 µm i.d. × 2 cm, packed with Acclaim PepMap100 C18, 5 µm, 100 Å (Thermo Scientific) for 8 min at flow 5ul/min, and then separated on a C18 reverse-phase column coupled to the Nano electrospray, EASY-spray (PepMap, 75mm x 50cm, Thermo Scientific) at flow 300 nl/min using an Dionex Nano-HPLC system (Thermo Scientific) coupled online to Orbitrap Mass spectrometer, Q Extactive Plus (Thermo Scientific). To separate the peptides, the column was applied with a linear gradient with a flow rate of 300 nl/min at 35 °C: from 1 to 35% in 100 min, from 35 to 55% in 43 min, from 55 to 90% in 5 min, and held at 90% for an additional 30 min, and then equilibrated at 1% for 20 min (solvent A is 0.1% formic acid, and solvent B is 80% acetonitrile, 0.1% formic acid). The Q Exactive was operated in a data-dependent mode. The survey scan range was set to 200 to 2000 m/z, with a resolution of 70,000 at m/z. Up to the 12 most abundant isotope patterns with a charge of ≥2 and less than 7 were subjected to higher-energy collisional dissociation with a normalized collision energy of 28, an isolation window of 1.5 m/z, and a resolution of 17,500 at m/z. To limit repeated sequencing, dynamic exclusion of sequenced peptides was set to 60 s. Thresholds for ion injection time and ion target value were set to 70 ms and 3 × 10^6^ for the survey scans and to 70 ms and 10^5^ for the MS/MS scans. Only ions with “peptide preferable” profile were analyzed for MS/MS. Data was acquired using Xcalibur software (Thermo Scientific). Column wash with 80% ACN for 40 min was carried out between each sample run to avoid potential carryover of the peptides.

### Data Analysis and Statistics of the protemomic data

For protein identification and quantification, we used MaxQuant software^34^, version 1.5.3.30. We used Andreomeda search incorporated into MaxQuant to search for MS/MS spectra against the UniProtKB database of Saccharomyces cerevisiae proteome, (Uniprot release, Aug 2016). The identification allowed two missed cleavages. Enzyme specificity was set to trypsin, allowing N-terminal to proline cleavage and up to two miscleavages. Peptides had to have a minimum length of seven amino acids to be considered for identification. Carbamidomethylation was set as a fixed modification, and methionine oxidation was set as a variable modification. A false discovery rate (FDR) of 0.05 was applied at the peptide and protein levels. An initial precursor mass deviation of up to 4.5 ppm and fragment mass deviation up to 20 ppm were allowed. Only proteins identified by more than 2 peptides were considered. To quantify changes in protein expression we used the label-free quantification (LFQ) using the MaxQuant default parameters^34^. For statistical and bioinformatic analysis, as well as for visualization, we used Perseus software (http://141.61.102.17/perseus_doku/doku.php?id=start). For functional enrichment analysis, the DAVID webserver^54^ was used. The STRING server (http://string-db.org/)^55^ was used to define protein interaction networks, which were visualized by using Cytoscape software^56^.

### RNA Purification for the transcriptomic analysis

Cells were incubated with Proteinase K (Epicentre MPRK092) and 1% SDS at 70°C to release the RNA. Cell debris was precipitated by centrifugation in the presence of KOAc precipitation solution. Finally, the RNA was purified from the supernatant using nucleic acid binding plates (UNIFILTER plates, catalog #7700-2810) and was stored with RNAse-inhibitor (Murine #M0314L) at −80°C.

### 3' RNA Library Preparation

Total RNA (˜20ng per sample) was incubated with oligo-dT RT primers with a 7bp barcode and a 8bp UMI (Unique Molecular Identifier) at 72oC for 3 minutes and transferred immediately to ice. RT reaction was performed with SmartScribe enzyme (TaKaRa Lot# 1604343A) at 42°C for one hour followed by incubation at 70°C for 15 minutes. Barcoded samples were then pooled and purified using SPRI beads X1.2 (AMPure XP). DNA-RNA molecules were tagmented using Tn5 transposase (loaded with oligo TCGTCGGCAGCGTCAGATGTGTATAAGAGACAG) and 0.2% SDS was used to strip off the Tn5 from the DNA, followed by a SPRI X2 clean up. NGS sequences were added to the tagmented DNA by PCR (KAPA HiFi HotStart ReadyMix 2X (KAPA Biosystems KM2605), 12 cycles). Finally, DNA was purified using X0.65 SPRI beads followed by X0.8 SPRI beads. The library was sequenced using Illumina NextSeq-500 sequencer.

### RNA Sequence analysis

Reads were mapped to the yeast genome (sacCer3) using bowtie2 with default parameters^57^. Duplicated reads were filtered using UMI, to remove PCR bias. To estimate the expression level of each gene we counted the number of reads that mapped to the 3' end of the gene (from 350bp upstream to 200bp downstream of TTS). The read counts in each sample were normalized to PPM (divided by the total number of reads and multiplied by 10^6^). The P-value of the differential expression analysis was obtained as described by Anders and Huber^58^, and corrected for multiple hypothesis testing using FDR.

### Imaging of the yeast cells

#### Microscopy used for Figure 2

Yeast strains were grown on synthetic media without uracil for selection. Imaging was performed using ScanR automated inverted fluorescent microscope system (Olympus). Images of cells were obtained in 384-well plates at 24°C using a 60× air lens (NA 0.9) with an ORCAER charge-coupled device camera (Hamamatsu). Images were acquired in the GFP channel (excitation filter 490/20 nm, emission filter 535/50 nm).

#### Confocal microscopy used for Figures 8- 9

The yeast cells were grown with anti-fluorescent medium, supplemented with casein and amino acids corresponding to plasmid selection and filtered with 0.22uM filters. Samples were grown overnight on anti-fluorescent medium diluted 1:10 and again 1:3 at OD_600_ of approximately 0.25. 500µl of sample was then placed on sterile µ-slides (Ibidi, GmbH, Munich, Germany) coated with Concanavalin A (C2272 SIGMA) and incubated several minutes before taking out excess. Cells were observed using time-lapse confocal microscopy (Olympus FV-1200) with 405nm and 488nm lasers. Images were taken every 20 minutes for 12 hours while at 30° and were analyzed using the ImageJ software.

### Scar counting and budding

Samples were collected after sorting by centrifuging at 3700g for 5 minutes; the pellet was resuspended with 500µl of PBS. After transferring liquid to Eppendorfs, samples were centrifuged at max speed for 1 minute and the pellet was resuspended with 4% paraformaldehyde and incubated 10 minutes. Paraformaldehyde was washed with PBS. The pellet was resuspended with 20µl of PBS and 2µl of Calcofluor (18909 SIGMA-ALDRICH) and incubated at room temp for 10 min. Samples were washed with PBS before placing on glass slides. Images were taken by confocal microscopy (Olympus FV-1200) and kept at 30^c^ throughout the process. Scars and buds were analyzed using the ImageJ software and manually counted.

## Acknowledgements

We are extremely grateful to the director of the confocal unit, Naomi Melamed-Book for assistance and running of the cell imaging assays. We thank Bruce Morgan for providing us with the roGFP expressing plasmids. We received financial support from the Marie-Curie integration grant (project number: 618806), the Israel Science Foundation (grant number: 1765/13), Human Frontier Science program (CDA00064/2014), and the US-Israel Binational Science Foundation (grant number: 2015056).

## Author contributions

MR, RF, OY, TR, NS, JG, NF, MS and DR designed the research and related experiments. MR, OY and WB established the FACS-based redox quantification. MR and WB performed yeast sorting. RF, MR, OY and DR performed proteomic analysis of the sorted yeast. JG performed the transcriptomic analysis of sorted cells. NS and MS designed the yeast strains expressing the roGFP sensor and performed imaging of the peroxisomal null strains. RF performed the cell imaging and scar detection, MR, RF and DR wrote the manuscript.

## Supplementary materials

### Figure legend

**Supplementary Figure 1.**
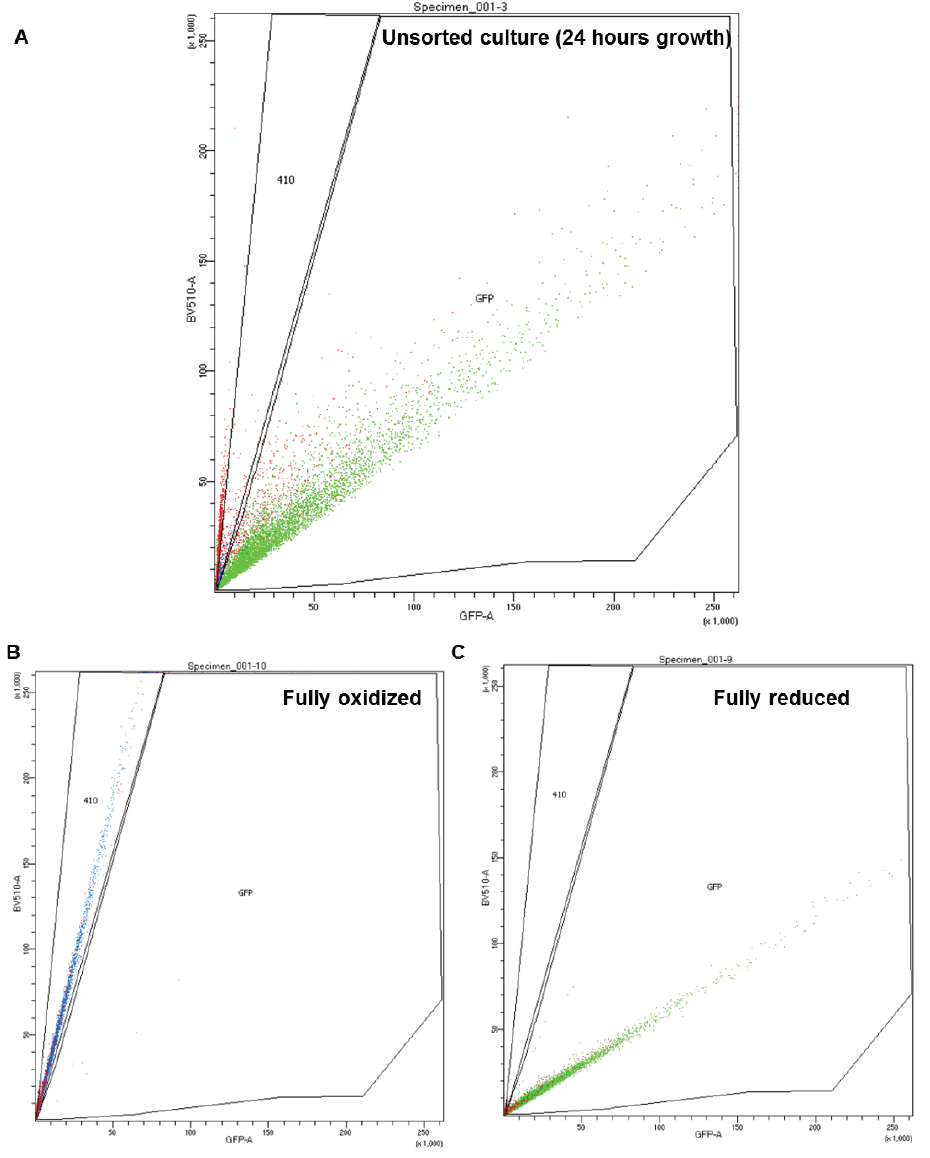
Representation of FACS subpopulation gates. (A) Representative data of an unsorted sample after 24h growth, plotted by the ratio between excitation at 405nm and 488nm (y and x axis, respectively). “GFP” corresponds with the “reduced” gate, while “410” corresponds with the oxidized. (B-C) Representative data of a fully oxidized or reduced sample (respectively).

**Supplementary Figure 2.**
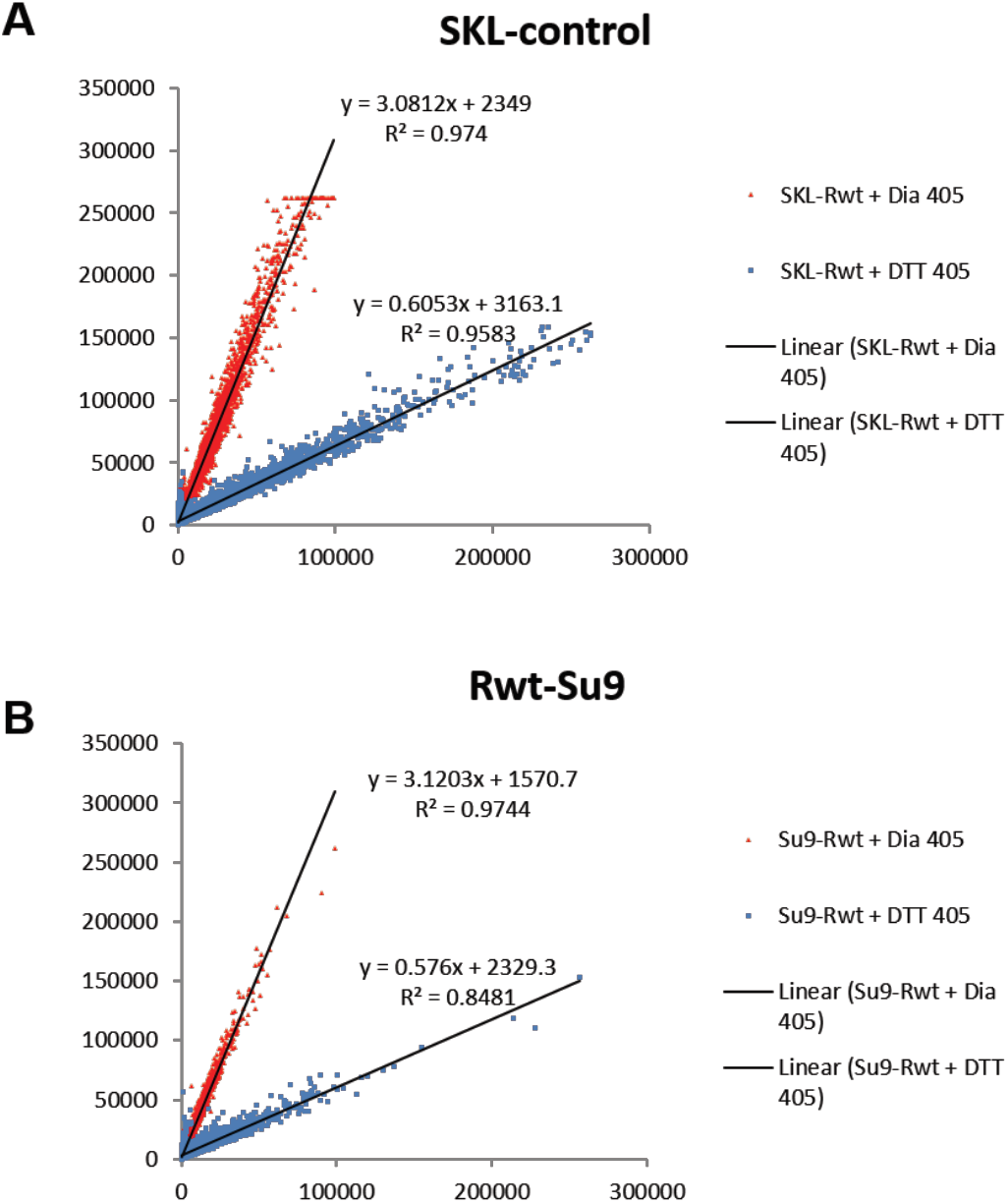
Linear characterization of (A) peroxisomal and (B) mitochondrial “oxidation gate”. Quantification of the redox status of fully reduced (blue) and fully oxidized (red) cells using FACS, using the peroxisomal and mitochondrial sensors.

**Supplementary Figure 3.**
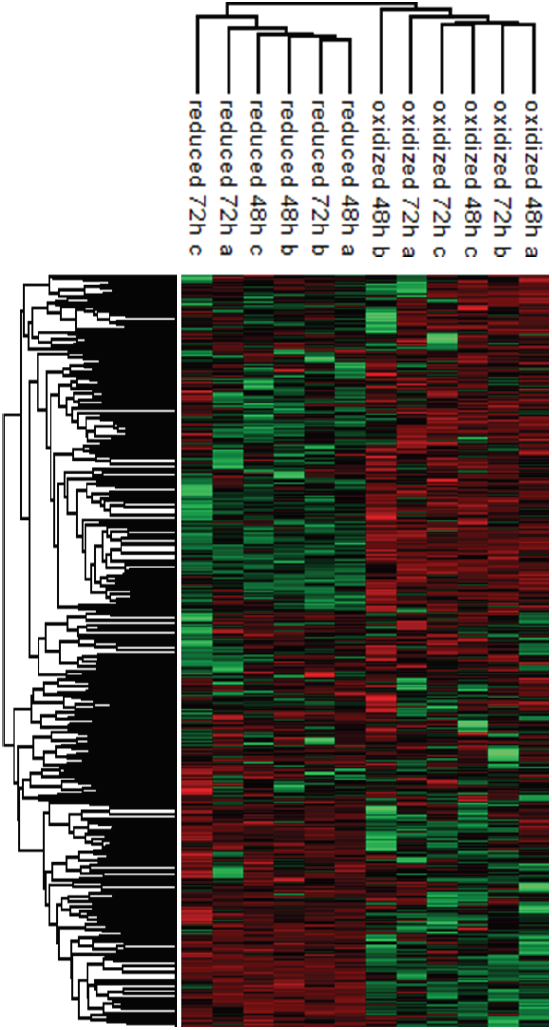
Hierarchical clustering of the protein abundances. log2 values of the log2 (LFQ) values of proteins identified in each post-sorting samples harvested at 48h and 72h (red corresponding to higher expression, green to lower). Median intensities were calculated and clustered in Fig 5A.

**Supplementary Figure 4.**
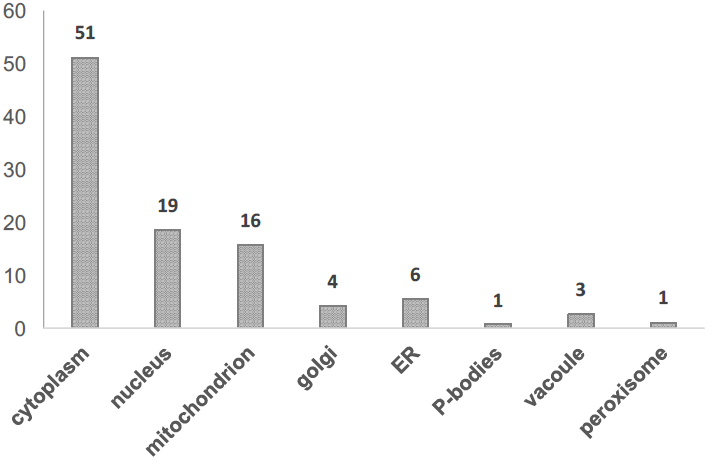
Cellular localization annotation of identified proteins. Annotation of all identified proteins, corresponding with the localization as agreed upon in the literature.

**Supplementary Figure 5.**
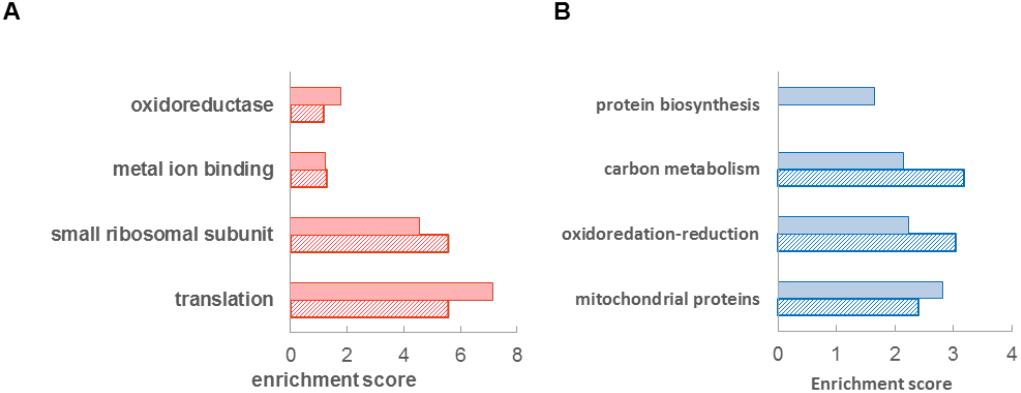
Functional enrichment of differentially expressed proteins (FDR <0.05) in the oxidized (red) and reduced (blue) subpopulations (corresponding to the volcano plot in Fig. 6).

**Supplementary Figure 6.**
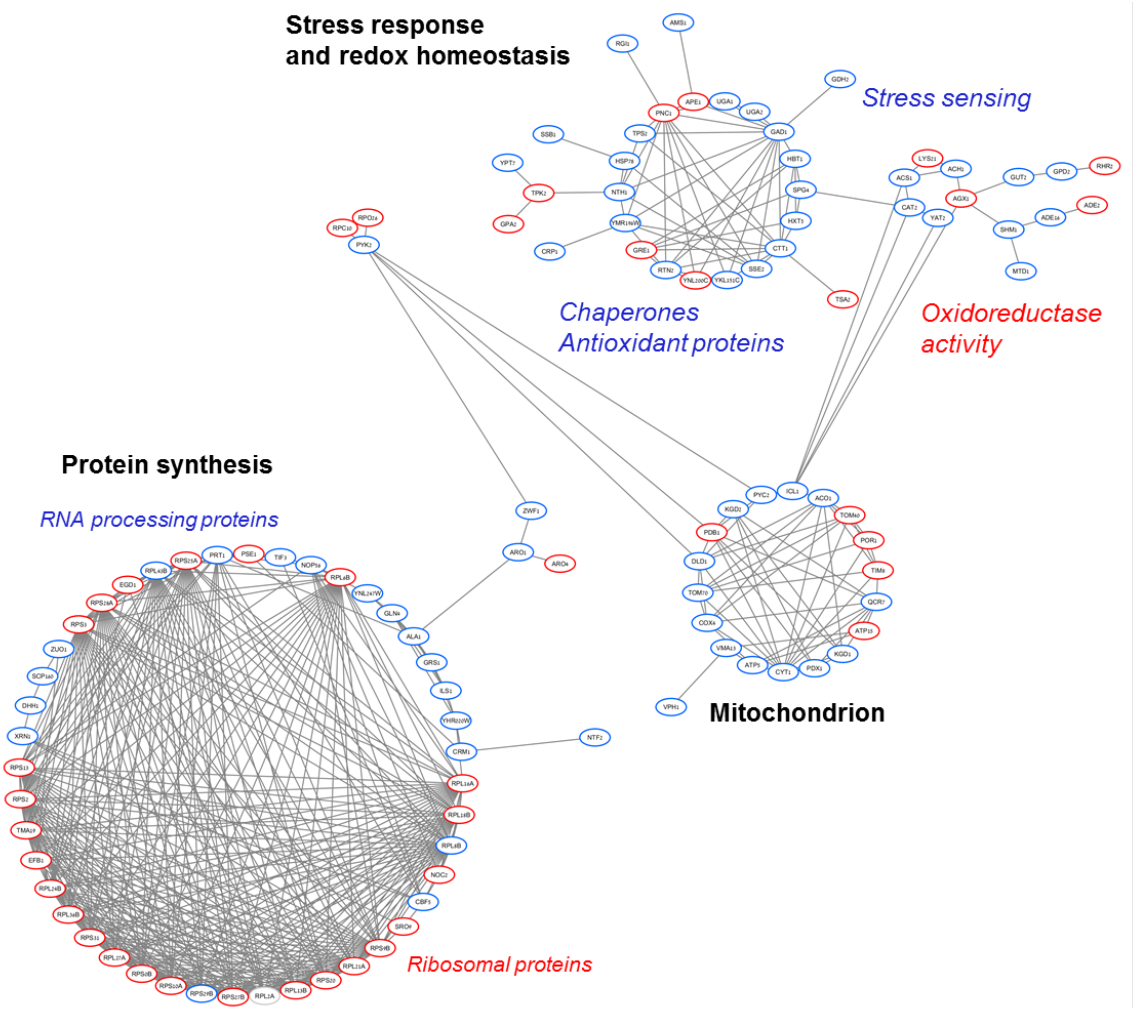
Interaction map of significantly differentially expressed protein identified in the 48h samples. Protein interactions are based on STRING database (as of January 2017) at highest confidence. Significantly upregulated proteins in the reduced subpopulation are in blue, upregulated proteins in the oxidized subpopulation are in red. Visualization was done by using Cytoscape.

**Supplementary Figure 7.**
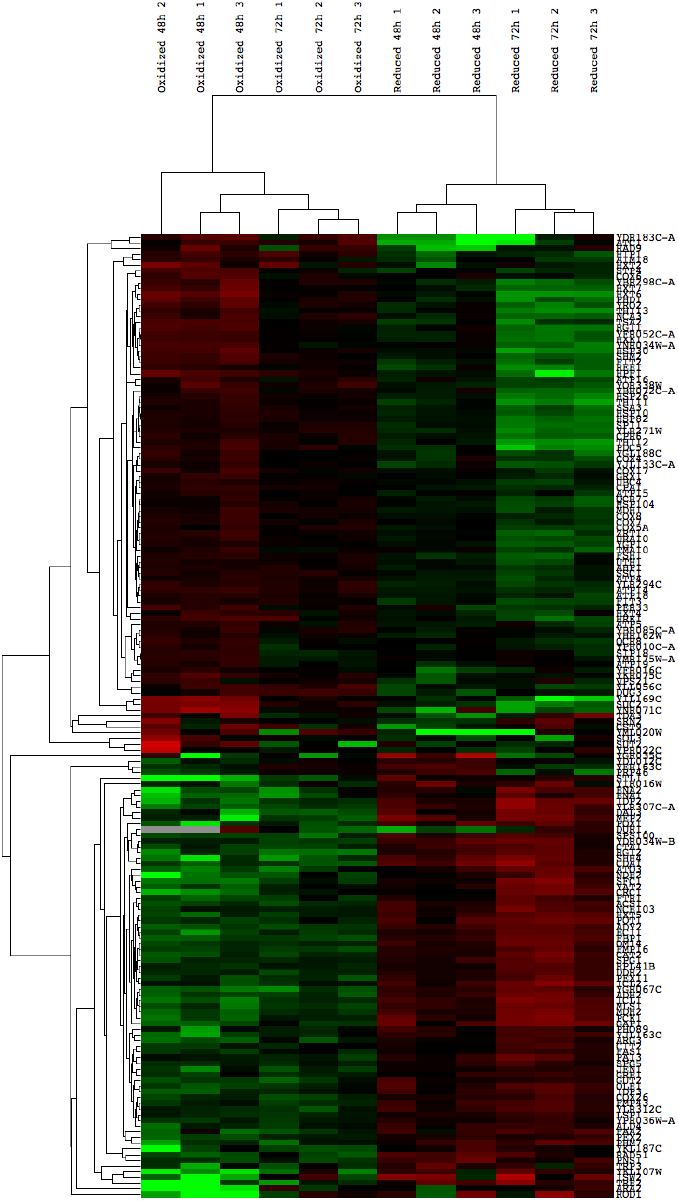
Hierarchical clustering of expression values of all differentially expressed genes (FDR<0.05) identified in the biological replicates of the post-sorting cells harvested at 48h and 72h. Transcripts above the median value of the row are in green, below in red. Each row was normalized by its median and log was taken for visualization purposes. The data was clustered using centered correlation similarity metric.

**Table S1.** Comparison of OxD values of wild type and knockout strains (related to Fig.2F).

| Strain | cytosol | peroxisome |
| --- | --- | --- |
| *Δpex3* | 0.90 | < 0.00001 |
| *Δcat2* | 0.25 | 0.001 |
| *Δatg36* | 0.16 | < 0.00001 |
| *Δahp1* | 0.40 | 0.003 |

The p-values for T-student test calculated using two tail similar variance.

**Table S2 (in excel)** - list of identified proteins in the proteomic analysis

**Table S3-4 (in excel)** significantly up-down regulated proteins in 48h and 72h growth.

**Table S5.** Comparison of wild type and knockout strains OxD values (related to Fig. 6E)

| Strain | Day 1 | Day2 | Day3 |
| --- | --- | --- | --- |
| *Δtsa2* | 1.27E-02 | 1.34E-04 | 2.60E-01 |
| *Δdhh1* | 1.36E-05 | 6.73E-08 | 1.86E-08 |
| *Δhbt1* | 5.76E-04 | 3.38E-05 | 5.04E-06 |
| *Δpnc1* | 3.50E-03 | 3.67E-05 | 3.05E-06 |
| *Δhsp30* | 4.51E-07 | 8.78E-08 | 8.32E-05 |
| *Δgdh2* | 6.37E-01 | 1.07E-05 | 4.63E-06 |

The p-values for T-student test calculated using two tail similar variance.

**Table S6 and S8** (**in excel)** – list of identified transcripts in the transcriptomic analysis

**Table S7-S9-11** (**in excel)** - significantly up-down regulated transcripts in 48h and 72h growth

**Table S12.**
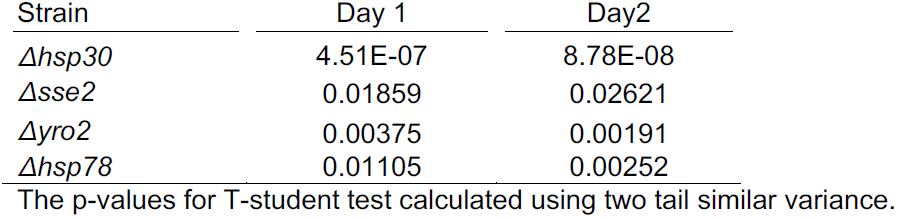
Comparison of wild type and knockout strains OxD values (related to Fig. 7)

